# CopySwitch - *in vivo* optimization of gene copy numbers for heterologous gene expression in *Bacillus subtilis*

**DOI:** 10.1101/446393

**Authors:** Florian Nadler, Felix Bracharz, Johannes Kabisch

## Abstract

The Gram-positive bacterium *Bacillus subtilis* has long been used as a host for production and secretion of industrially relevant enzymes like amylases and proteases. It is imperative for optimal efficiency, to balance protein yield and correct folding. Gene copy numbers are an important tuning valve for the optimization of heterologous gene expression. While some genes are best expressed from many gene copies, for other genes, medium or even single copy numbers are the only way to avoid formation of inclusion bodies, toxic gene dosage effects or achieve desired levels for metabolic engineering. In order to provide a simple and robust method to address above-mentioned issues in the Gram-positive bacterium *Bacillus subtilis*, we have developed an automatable system for the tuning of heterologous gene expression based on the host’s intrinsic natural competence and homologous recombination capabilities. By supplying our reporter strains with a linearized, low copy number plasmid containing homology regions left and right of the reporter genes and an antibiotic resistance marker, we could show an up to 3.6-fold higher *gfp* (green fluorescent protein) expression and up to 1.3-fold higher *mPLC* (mature phospholipase C) expression after successful recombination and thus circularization of our plasmid. Furthermore, the plasmid-borne *gfp* expression seems to be more stable, since over the whole cultivation period the share of fluorescent cells compared to all measured cells is consistently higher.

## 2. Introduction

Heterologous gene and pathway expression remains a task mostly approached by trial and error, making use of a broad palette of available optimization strategies (Stevens, 2000). The goal of any approaches is to find the optimal level of gene expression for the maximum amount of active protein without the formation of inclusion bodies. In prokaryotic gene expression this implies finding a suited balance between transcription and translation speed on the one hand and correct protein folding on the other hand. This can be achieved by the simple measure of reducing the temperature during protein production with the drawback of increased process times (Sørensen and Mortensen, 2005).

Applying molecular tools that have been developed to easily find optimum conditions are very helpful. For this, modern DNA assembly techniques are used to create a library of constructs, transcribed and translated with different strengths, for example by varying the promoter (Kraft et al., 2007), the ribosome binding site (Kohl et al., 2018) or by fusion with peptides aiding in correct folding (Butt et al., 2005; Kraft et al., 2007). Another approach is to vary the copy number of the genetic constructs. As shown before, the number in which a gene is present in a cell is correlated with the expression level of said gene (Lee et al., 2015). In the most prominent host *Escherichia coli*, plasmids with different copy numbers are available such as the high copy number pUC vectors (Yanisch-Perron et al., 1985) and the low copy number pBR322 based vectors (Bolivar et al., 1977) and assisted integration into the chromosome is possible through recombineering (Sharan et al., 2009). Plasmid replication is a major burden on the cell’s metabolism, thus always requiring an active selection mechanism to ensure plasmid stability during production. The higher the copy number of the plasmids, the higher this burden and the less stable the plasmid DNA is propagated. For the yeast *S. cerevisiae* it was demonstrated, that a set of tunable copy number plasmids (dependent on antibiotic concentration) could balance a pathway for n-butanol production in a way that yielded a 100-fold increase in production of the desired chemical (Lian et al., 2016). In another publication, different copy number plasmids were introduced into *Bacillus methanolicus* via electroporation to show the effect of gene dosage on the expression of *gfpuv*, an amylase from *Streptomyces griseus* and the lysine decarboxylase *cadA* from *E. coli* for overproduction of cadaverine. A positive correlation between estimated plasmid copy numbers and observed expression levels could be determined (Irla et al., 2016). Since plasmid copy numbers can be affected by media composition as well as growth phase, efforts have been undertaken to uncouple expression levels from gene dosage by introducing so-called incoherent feedforward loops in the form of transcription-activator-like effector (TALE)-regulated promoters (Segall-Shapiro et al., 2018).

Contrary to the Gram-negative organism *E. coli*, which under laboratory conditions is only able to take up plasmid DNA after various pretreatment steps, *Bacillus subtilis* gains natural competence under certain circumstances, which makes transforming *B. subtilis* as easy as cultivating the cells in a competence-inducing medium in the presence of the DNA to be integrated (e.g. plasmid DNA) for a few hours (Kumpfmüller et al., 2013). To further accelerate this process, *B. subtilis* strains with inducible competency have been developed (Rahmer et al., 2015). In contrast to this easy method for transforming linear DNA for genomic integration, the transformation of circular plasmid DNA is more difficult in *B. subtilis*. One method based on naturally competent cells has the drawback of needing multimeric plasmid forms (Canosi et al., 1978), which have to be generated either by time-consuming and potentially error-prone PCR, followed by ligation or by transformation of a monomeric form into specialized *E. coli* strains (e.g. JM105 (Voss et al., 2003)), yielding partly multimeric plasmid fractions. Most other methods of transforming *B. subtilis* with replicative plasmids require the use of special devices (e.g. electroporation), pretreatment of cells with polyethylene glycol (PEG) or glycine or the preparation of protoplasts, which is tedious and requires recovery for 2 - 3 days (Vojcic et al., 2012).

The CopySwitch system presented in this work allows the rapid and easy *in vivo* increase of copy numbers for applications in *B. subtilis* without the drawbacks of classic plasmid transformation techniques. The underlying process is depicted in Figure 1. A construct for heterologous expression is integrated into the chromosome via natural competence. After verification of the correct insertion, a linearized plasmid, containing an origin of replication (*ori*) for *B. subtilis*, an antibiotic resistance gene and homologous regions up- and downstream of the integration site, is added to competent cells. Upon intracellular uptake, recombination occurs and all genetic elements between the two recombination sites are copied onto the plasmid, rendering it circular and therefore replicative again (Tomita et al., 2004b).

**Figure 1.**
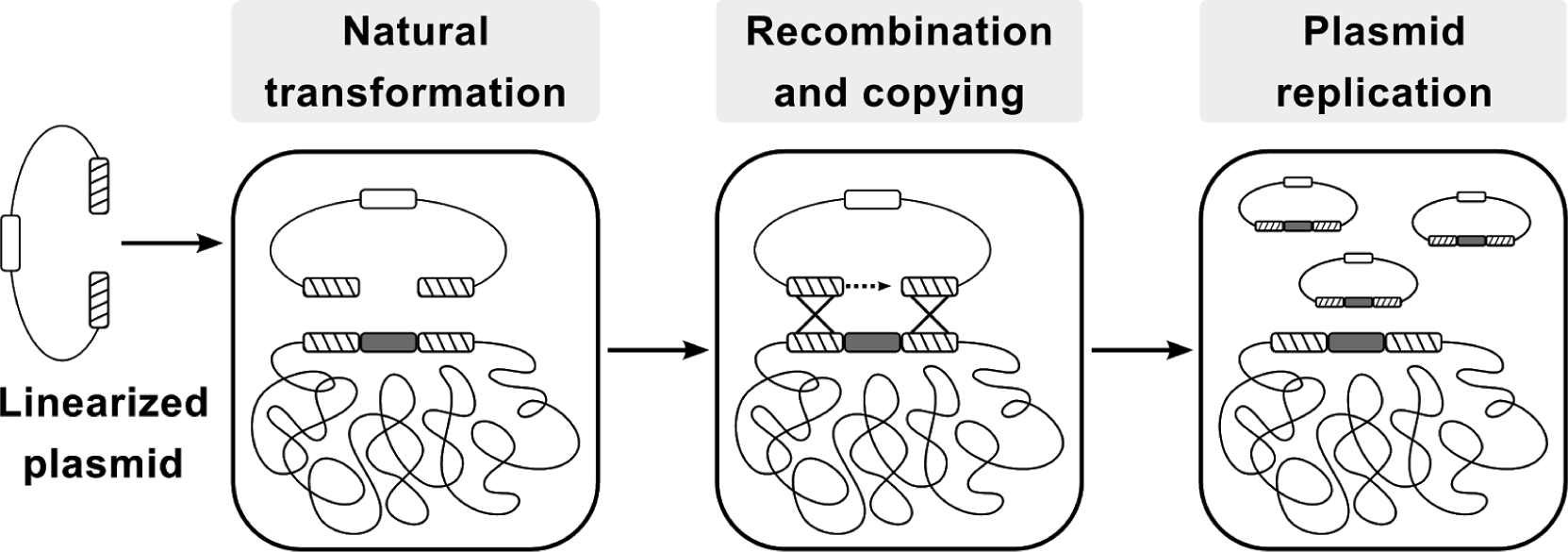
Flow-chart of the CopySwitch method; after linearization of a replicative plasmid (via restriction or PCR), it is taken up by *Bacillus subtilis* via natural transformation. Regions homologous to the chromosome are recombined and the DNA sequence between the regions is copied onto the plasmid. Hatched blocks: homology regions; dark grey blocks: sequence of interest; white block: low copy number *ori*; Adapted from Tsuge and Itaya (Tsuge and Itaya, 2001).

We chose two different reporter systems for demonstrating the application of CopySwitch, one being the intracellular production of GFP and the other being the secretion of a recombinant mature phospholipase C (mPLC) from *Bacillus cereus* SBUG 516, both under the acetoin-inducible promoter *P_acoA_* from *B. subtilis*. The mPLC is a monomeric, phosphatidylcholine-preferring PLC (E.C.3.1.4.3, UniProt-ID P09598), containing three catalytically relevant zinc ions. Its sequence comprises of the amino acids 39-283 (mature form) and is preceded by the first 33 amino acids of the *B. subtilis amyE* gene, which function as a signal peptide for secretion (Durban et al., 2007). Green fluorescent protein production can be assessed via flow cytometry fluorescence measurements, whereas mPLC activity is evaluated by the increased absorbance at 410 nm due to accumulation of *p*-nitrophenol from the cleavage of the chromogenic substrate *p*-NPPC (*para*-nitrophenylphosphorylcholine) (Durban et al., 2007; Kurioka and Matsuda, 1976).

With the here described method we aim to address some of the shortcomings and current bottlenecks of traditional replicative plasmid transformation protocols.

## 3. Materials and Methods

### 3.1 Strains and plasmids

For propagation and maintenance of plasmids *E. coli* DH10B cells were used. For all experiments on *gfp* expression, a derivative of *Bacillus subtilis* 6051-HGW was used as a starting point, combining six different modifications previously described (Kabisch et al., 2013; Zobel et al., 2015) (see Table 2). For all experiments on *mPLC* (matured phospholipase C) expression, a derivative of *Bacillus subtilis* PY79 was used, combining seven knock-outs of extracellular proteases. This so-called KO7 strain is available from the Bacillus Genetic Stock Center (Columbus, OH, USA) under the BGSCID 1A1133. In our studies, we were using a derivative of pBS72 (Titok et al., 2003) for a low copy number *ori*. Instead of the full length *B. subtilis* origin region as it is used for example in the commercially available pHT01 system (MoBiTec, Göttingen, Germany), we shortened the region by approximately 600 base pairs (bps) by deleting two putative open reading frames (‘orf-4 and orf-3) of unknown function (Titok et al 2006) to reduce replicative burden.

For details on the plasmid sequences including oligonucleotides see Supplemental Material section.

### 3.2 Media and cultivation

All standard cultivations were performed in LB containing 10 g/L NaCl (LB10; Carl Roth, Karlsruhe, Germany), except for *B. subtilis* strains carrying a resistance gene for the salt-sensitive antibiotic zeocin (InvivoGen, San Diego, CA, USA). In these cases, LB containing only 5 g/L NaCl (LB5; Carl Roth, Karlsruhe, Germany) was used. Transformation medium for *Bacillus subtilis* was prepared according to Kumpfmüller (Kumpfmüller et al., 2013). Antibiotics were used in the following concentrations: 100 μg/ml ampicillin, 50 μg/ml (*E. coli*) or 12.5 - 25.0 μg/ml (*B. subtilis*) kanamycin, 100 μg/ml spectinomycin and 40 μg/ml zeocin.

All incubations for the flow cytometry experiment were performed at 37°C and 200 - 250 rpm in a 25 mm amplitude rotary shaker (Infors GmbH, Einsbach, Germany). Thirty milliliters of preculture (15 ml 2x LB5, 1% (v/v) glucose from a 40% (w/v) stock solution, 40 μg/ml zeocin and with or without 25 μg/ml kanamycin, filled to 30 ml with sterile, desalted H_2_O) were inoculated from glycerol stocks of BsFLN045 and BsFLN051, respectively. After incubation overnight, 20 ml of main culture (10 ml 2x LB5, 0.5% (v/v) glucose from a 40% (w/v) stock solution, 0.5% (v/v) acetoin from a 10% (w/v) stock solution, 40 μg/ml zeocin and with or without 25 μg/ml kanamycin, filled to 20 ml with sterile, desalted H_2_O) were inoculated with preculture to an optical density of 0.1 at 600 nm (OD_600_) in quadruplicates for each strain. Media were prepared as master mixes and distributed evenly into 50 ml Erlenmeyer flasks for cultivation.

Cultivations for the phospholipase C experiments were performed at 30°C and 150 rpm in a 50 mm amplitude rotary shaker (Eppendorf New Brunswick, Enfield, CT, USA). Twenty milliliters of preculture (SB medium: 32.0 g/L Tryptone, 20.0 g/L yeast extract, 5.0 g/L NaCl, 5.0 ml 1N NaOH, 42.3 mM Na_2_HPO_4_, 22.0 mM KH_2_PO_4_, 18.7 mM NH_4_Cl and 0.02 mM CaCl_2_, supplemented with 1% (v/v) glucose from a 40% (w/v) stock solution, 40 μg/ml zeocin and with or without 25 μg/ml kanamycin) were inoculated from glycerol stocks of BsMKA005, BsFLN064 and BsFLN066, respectively. To ensure sufficient growth overnight, precultures were incubated at 37°C. Fifty milliliters of main culture (SB containing 0.5% (v/v) of acetoin from a 10% stock solution in SB and respective antibiotic) were inoculated to reach an OD_600_ of 0.2 in triplicates for each strain. Cultivation was carried out in 500 ml Erlenmeyer flasks.

### 3.3 Enzymes

All restriction enzymes and Q5 proofreading polymerase (for amplicons > 3 kb) were purchased from New England Biolabs (NEB, Ipswich, MA, USA). Colony polymerase chain reactions (cPCRs) were performed with standard Taq polymerase, whereas error-free amplification of small (< 3 kb) fragments was performed with OptiTaq, which is a mixture of Taq and Pfu polymerase (Roboklon, Berlin, Germany). Alternatively to Q5, proofreading Polymerase X was also used for amplicons > 3 kb (Roboklon, Berlin, Germany).

### 3.4 Molecular biology methods

All PCRs and restriction digestions were carried out as suggested by the supplier of the used enzymes. Sequencing reactions and oligonucleotides were purchased from Eurofins genomics (Ebersberg, Germany). If replicative plasmids were used as PCR templates and no antibiotic switch was possible after cloning, a thorough DpnI digestion (1 - 2 hours at 37°C) was performed to reduce background through template plasmid carry-over, followed by heat inactivation (20 min at 65°C). PCR fragments were column-purified before being used for cloning (NucleoSpin^®^ Gel and PCR Clean-up Kit, Macherey-Nagel, Düren, Germany or Monarch^®^ PCR & DNA Cleanup Kit, NEB, Ipswich, MA, USA). Plasmids were also column-purified with two different kits (High Pure Plasmid Isolation Kit, F. Hoffmann-La Roche AG, Basel, Switzerland or Monarch^®^ Plasmid Miniprep Kit, NEB, Ipswich, MA, USA). For plasmid preparation from *B. subtilis*, 5 - 10 mg of lyophilized lysozyme (Sigma-Aldrich, St. Louis, MO, USA) were added to the resuspension buffer and incubated at 37°C for 10 min. All subsequent steps were done as described in the respective protocol. DNA assemblies were performed using SLiCE as described by Messerschmidt and coworkers (Messerschmidt et al., 2016). In contrast to the described method, DNA was transformed by electroporation (2.5 kV, 2 mm gap). For an assembly containing more than three DNA parts, amplifying the SLiCE assembled DNA via PCR enabled us to reduce the fragment number. Correct assembly of fragments and integration into a *B. subtilis* genome was detected by colony PCR (cPCR) and confirmed by sequencing of either plasmids or proofreading amplified PCR products of regions of interest.

### 3.5 Transformation of *Bacillus subtilis*

For transformation, 0.05% (w/v) of casamino acids (CAA; Sigma-Aldrich, St. Louis, MO, USA) and either 1% (v/v) glycerol stock or 3 - 5% (v/v) of an overnight culture of the desired strain (grown in the same medium) were added to transformation medium (TM) containing a suitable antibiotic. Cells were cultivated for 5 - 6 hours, shaking at 37°C until the broth was visibly turbid. At this point, plasmid DNA (undigested for chromosomal integration and *XhoI*-linearized for CopySwitch) was added at a final concentration of > 200 fmol/ml and incubated for 30 minutes while shaking. For each milliliter of transformation sample, 200 μl of expression mix (2.5% (v/v) of yeast extract and CAA each) were added and, depending on the task, shaken for another one (chromosomal integration) to four (CopySwitch) hours at 37°C. Afterwards, different amounts (200 to 1000 μl) were spread onto antibiotic-containing LB-agar-plates and incubated overnight to ensure growth of distinct single colonies. To recycle the antibiotic marker in the chromosomal integration approaches, single colonies were subjected to xylose-induced expression of *cre* recombinase by growth in 300 μl of LB5-Zeo (40 μg/ml) containing 1% (v/v) of xylose for > 6 hours at 37°C and 1400 rpm (Thermoshaker, Eppendorf, Hamburg, Germany). Constant aeration during this time was achieved by melting a hole into the lid of the 2 ml Eppendorf tubes used. This procedure results in recombination of *lox71* and *lox66*, thereby looping out the marker gene cassette between and generating a *lox72* scar. After a dilution streaking on zeocin-containing LB5-agar-plates, single colonies were further verified via cPCR, replica-plating to ensure loss of the antibiotic marker and finally sequencing. To verify a CopySwitch, we isolated plasmid from *B. subtilis* and transformed the obtained DNA into *E. coli* to obtain sufficient amounts of plasmid for sequencing.

### 3.6 Enzymatic assay for mPLC activity

#### 3.6.1 Sampling

Samples for the enzymatic assay of mPLC activity were collected after 6, 12, 24 and 48 hours of cultivation. Two milliliters of culture broth were centrifuged (17.000 rcf, 4°C, 5 min) and 1.8 ml were transferred into new vessels. Sterile-filtering was omitted to ensure full enzymatic activity.

#### 3.6.2 Assay principle and plate reader measurements

The underlying principle of the activity assay lies in the ability of mPLC to turn the non-natural substrate *p*-NPPC (CAS number 21064-69-7, Cayman Chemicals, Ann Arbor, MI, USA) into phosphorylcholine and *p*-nitrophenol. Absorption of *p*-nitrophenol is measured at 410 nm (Durban et al., 2007; Kurioka and Matsuda, 1976). Activity measurements were carried out with 70 μl of diluted supernatant (one part supernatant + three parts 100 mM borax-HCl buffer, pH 7.5) and 30 μl of substrate solution (100 mM *p*-NPPC dissolved in 100 mM borax-HCl buffer, pH 7.5) in 96-well microtiter plates (96-well, PS, F-bottom, #655101, Greiner Bio-One GmbH, Frickenhausen, Germany). Measurements were done at 30°C for two hours every minute and for another five hours every five minutes afterwards. Since substrate depletion caused the reaction to plateau after several hours, only values from the first four hours were taken for all calculations.

#### 3.6.3 Data Analysis

All data conversion, analyses and plotting was performed with R Version 3.5.1 (Ihaka and Gentleman, 1996) using Wickhams tidyverse Version 1.2.1. (Wickham, 2017)

### 3.7 Assaying GFP fluorescence

#### 3.7.1 Sampling

Samples for flow cytometry analysis were taken after 3, 5, 7, 9, 11, 20 and 24 hours of cultivation, to ensure capturing the transition between non-expressing and gene-expressing state. The sample volume was chosen to yield an OD_600_ between 1.7 and 2.6 after pelleting the cells, discarding the supernatant and resuspending in 1.5 ml 1x ClearSort Sheath Fluid (Sony Biotechnology, Weybridge, UK). Before resuspension, the cell pellets were stored at -20°C until all samples were collected and prepared for measurement. Reconstituted samples were diluted again (one part sample plus four parts 1x ClearSort Sheath Fluid), to be in a suitable range of events per second for flow cytometry measurement.

#### 3.7.2 Flow Cytometry

Fluorescence in cells was detected in a Sony LE-SH800SZBCPL (Sony Biotechnology, Weybridge, UK) using a 488 nm argon laser. Photomultipliers for backscatter and FL-1 (525/50 nm) were set to 25.5 % and 38.0 % with a FSC-threshold of 0.20 % and a window extension of 50. The FSC-diode was set on an amplification setting of 6/12 with events per second being 50’000.

#### 3.7.3 Flow Cytometry Data Analysis

Areas of scattering and fluorescence signals (FSC-A, SSC-A, FL1-A) were brought to a near-normal distribution by applying an inverse hyperbolic sine (asinh) to all values. Cell agglomerates and fragments were excluded by fitting a bivariate normal distribution in the channels FSC-A and FSC-W. For the threshold between fluorescent and non-fluorescent cells, the minimum of the clearly bimodal distribution of all measured FL-1 values was chosen. For every sample, medians of the fluorescent population as well as the ratio of counts between fluorescent events and total events was calculated. The resulting values were analyzed by showing means and standard deviations of biological quadruplicates. All analyses, calculations and plotting were performed with R (Ihaka and Gentleman, 1996). Specifically, for parsing of fcs files and gating, bioconductor’s (Huber et al., 2015) flowcore (Hahne et al., 2009) was used. Packages from the metapackage tidyverse, especially dplyr (Hadley Wickham, Romain Francois, Lionel Henry and Kirill Müller, 2017) and ggplot2 (Wickham, 2011) were employed for summarization and presentation.

#### 3.7.4 Plate reader measurements

Directly after all flow cytometry measurements had been done, 200 μl of each diluted sample were transferred from the flow cytometry tubes into 96-well microtiter plates (Nunc™ MicroWell™ #152037, ThermoFisher Scientific, Waltham, MA, USA). Optical density was measured at 600 nm in a PHERAstar FSX (BMG Labtech, Ortenberg, Germany). Fluorescence was induced with an excitation wavelength of 485 nm and measured at an emission wavelength of 520 nm. This procedure guarantees comparability between single-cell- and population-based data. All analyses, calculations and plotting were performed with R (Ihaka and Gentleman, 1996).

## 4. Results

### 4.1 Optimizing parameters for successful transformation and gene transfer onto replicative plasmid

Transformation via natural competence requires internalization of single-stranded DNA, formation of a heteroduplex between regions of homology and expression of an antibiotic resistance marker. Using our method for replicative plasmid transformation furthermore requires time for expression of additional, plasmid-based replication machinery (e.g. RepA protein) and replication of the newly circularized plasmid. Three different parameters to optimize the yield of plasmid-bearing colonies were tested. For standard chromosomal integration we used homology regions of 500 bp and a regeneration time after transformation of one hour. However, this procedure did not yield any colonies for CopySwitch, regardless of the antibiotic concentration (see Table 1). To increase the probability of correctly assembled replicative plasmids, longer regeneration times (one, two and four hours) and longer regions of homology (500, 1000 and 2000 bp) were tested. The volume plated corresponds to 48 fmol of linearized plasmid in the transformation mix. All of the different conditions were tested in BsFLN045 with either plasmids containing different homology regions or elution buffer (negative control). From the colony count of these transformations, the conclusions can be made that i) four hours regeneration time resulted in satisfactory CopySwitch efficiency and ii) 2000 bp are a sufficient length for homology regions. Further, a kanamycin concentration of 25 μg/ml was superior to both 12.5 μg/ml (with the latter, a lot of false-positive clones arose after > 17 h incubation period, data not shown) and 50 μg/ml (too high, no colonies formed at all). To confirm that the higher colony count was not only due to replication of correct clones because of elongated regeneration times, the OD_600_, which is correlated to the number of cells in suspension, was measured directly before plating. In the case of 2000 bp and 12.5 μg/ml, the OD for example was only doubled between one and four hours of regeneration time, whereas the colony forming units (cfu) differed by a factor of 35 for these two time points. Sequencing of three different, re-transformed plasmids showed no mutations of the copied locus.

**Table 1:** Optimization of transformation conditions

| Regeneration time after transformation | 1 h |  |  | OD<sub>600</sub> | 2 h |  |  | OD<sub>600</sub> | 4 h |  |  | OD<sub>600</sub> |
| --- | --- | --- | --- | --- | --- | --- | --- | --- | --- | --- | --- | --- |
| Kanamycin [µg/ml] | 12.5 | 25 | 50 |  | 12.5 | 25 | 50 |  | 12.5 | 25 | 50 |  |
| BsFLN049 (500 bp) | 0 | 0 | 0 | 3,2 | 0 | 0 | 0 | 4,6 | 0 | 0 | 0 | 6,2 |
| BsFLN050 (1000 bp) | 0 | 0 | 0 | 3,2 | 0 | 0 | 0 | 4,6 | 0 | 0 | 0 | 6,1 |
| BsFLN051 (2000 bp) | 1 | 0 | 0 | 3,2 | 9 | 0 | 0 | 4,8 | 35 | 25 | 0 | 6,4 |
| Neg. control (elution buffer) | 0 | 0 | 0 | 3,0 | 0 | 0 | 0 | 5,2 | 1 | 0 | 0 | 6,0 |

**Table 2:**
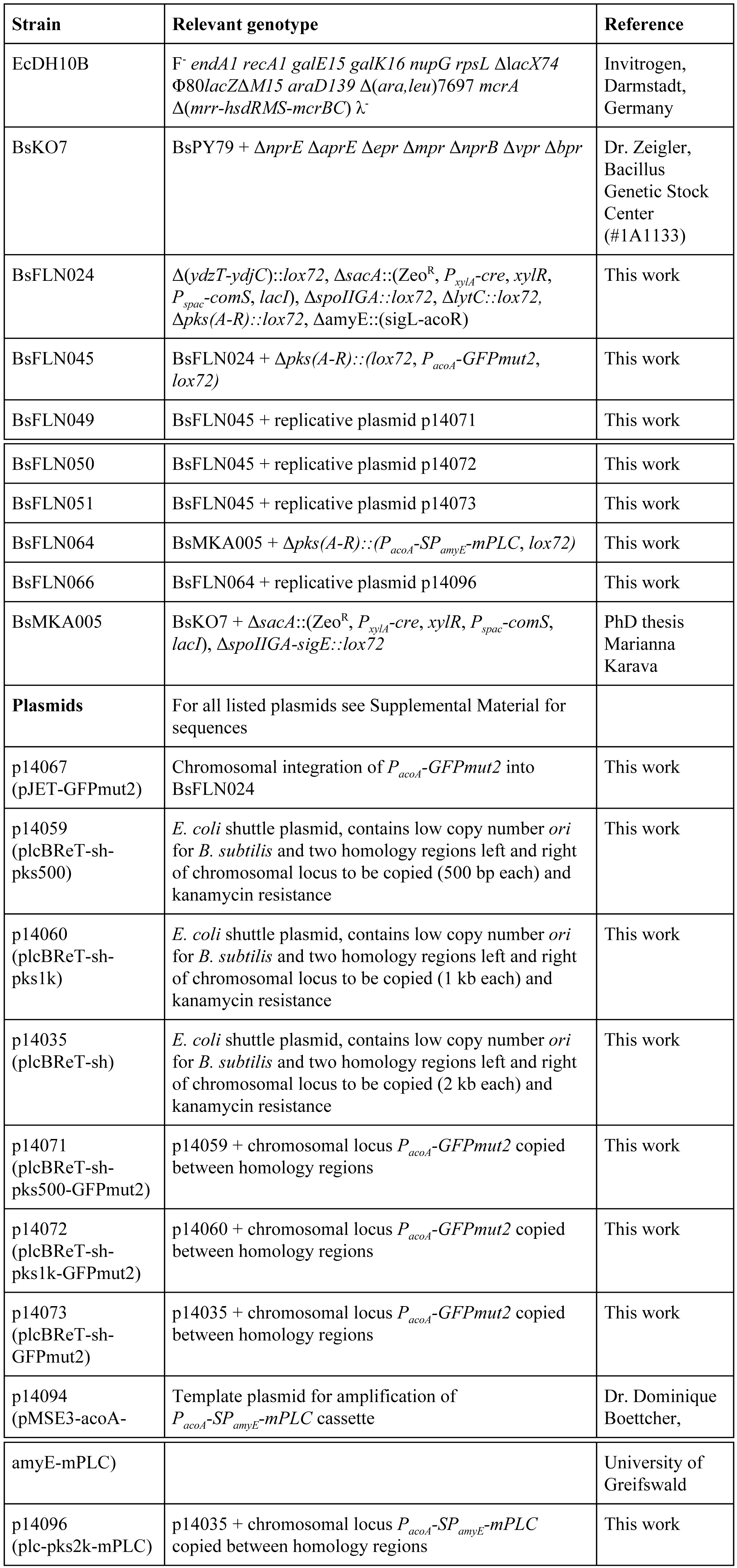
Strains and plasmids used in this work.

### 4.2 Growth curve comparison between one and multiple gene copy bearing strains

The averaged growth curves of a strain containing only a chromosomal copy of *gfp* (BsFLN045) and a strain containing a chromosomal and a plasmid copy (BsFLN051), all under the control of an acetoin-inducible promoter (*P_acoA_*), have been recorded at 37°C. During lag- and log-phase there is no visible difference between both curves. Contrary to this observation, in transient phase the plasmid-bearing strain is growing slower and does not reach an OD_600_ as high as BsFLN045 (see Figure 2A).

**Figure 2:**
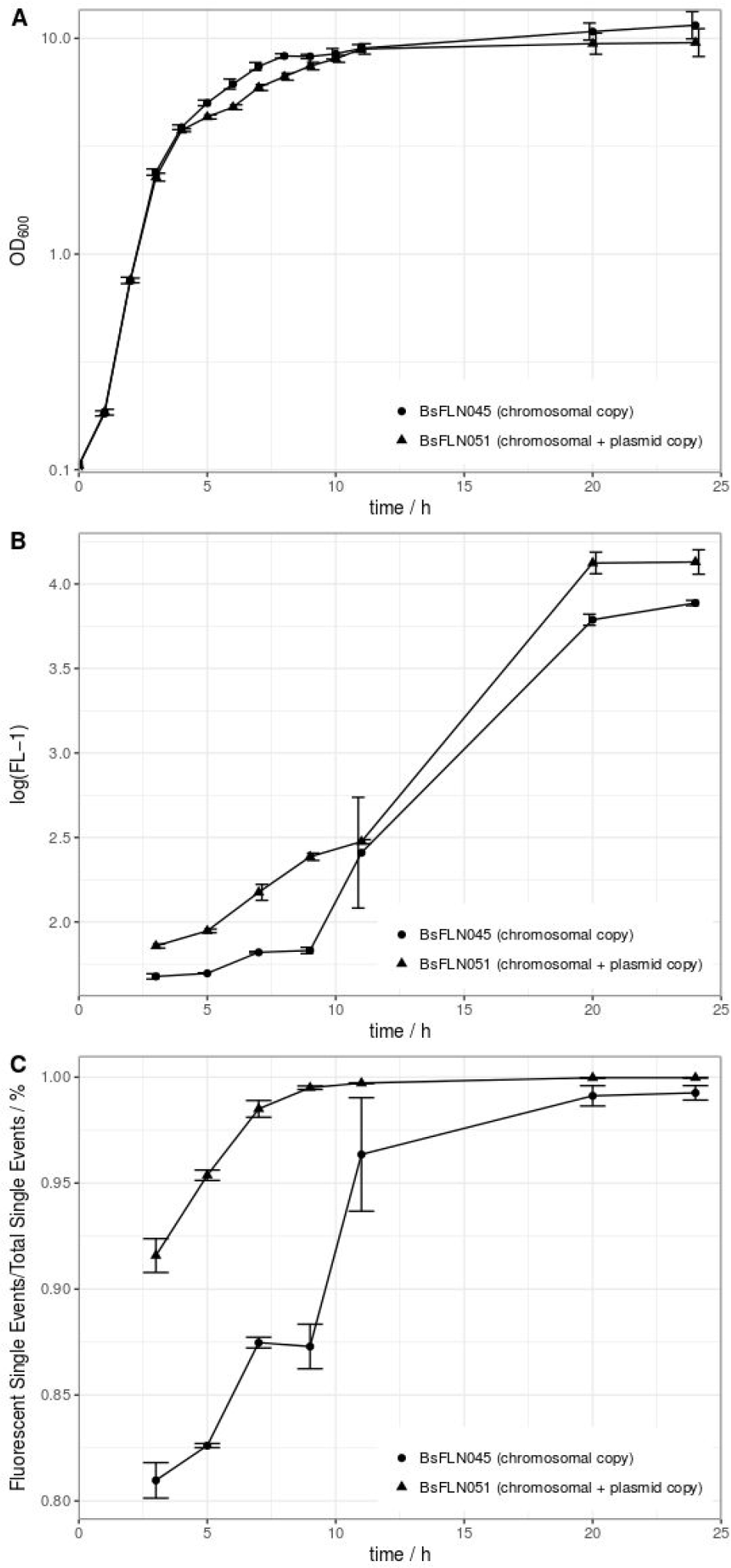
**A** Averaged growth curves of strains BsFLN045 (chromosomal copy) and BsFLN051 (chromosomal + plasmid copy); **B** Logarithmic plot of increase in FL-1 fluorescence and assumed intracellular GFP content over cultivation time. For all points except at t = 11 h, the increased fluorescence of BsFLN051 is statistically significant (Student’s t-test, alpha = 0.05). **C** Increase in the share of *gfp*-expressing cells over cultivation time. For all points except at t = 11 h the increased fluorescence of BsFLN051 is statistically significant (Student’s t-test, alpha = 0.05). Plotted values are averages of n = 4 biological replicates with error bars showing standard deviation.

All strains of the mPLC experiment were also investigated for differences in growth behaviour. At 30°C, no significant differences could be found (see Figure 3A). The reason for choosing 30°C instead of 37°C for cultivation, lies in higher protein stability and solubility during heterologous expression at lower temperature (Schein and Noteborn, 1988).

**Figure 3:**
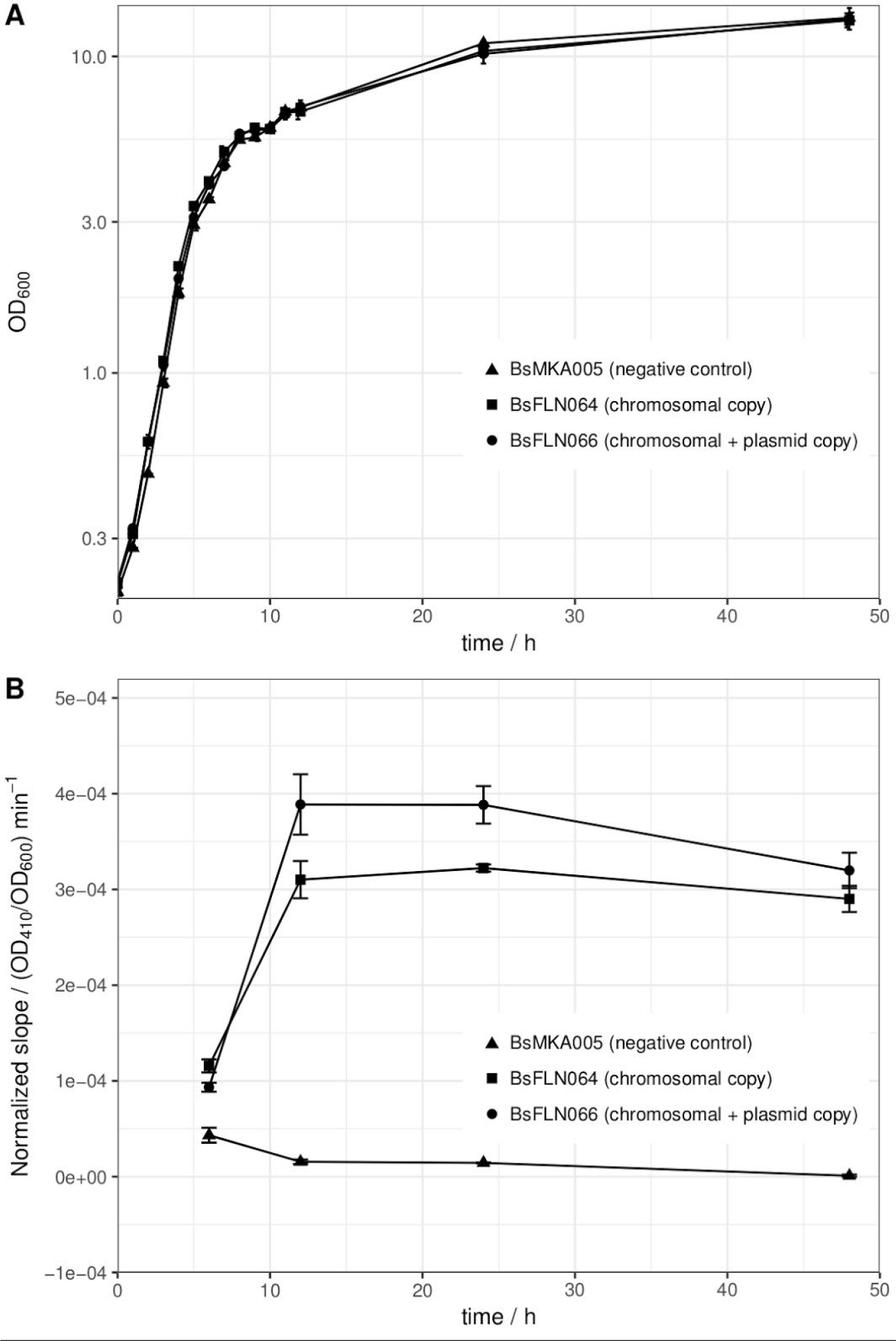
**A** Averaged growth curves of strains BsMKA005 (negative control), BsFLN064 (chromosomal copy) and BsFLN066 (chromosomal + plasmid copy); **B** OD_600_-normalized slope values of phospholipase C assay as a measure of enzymatic activity. Since amount of substrate and volume of enzyme solution (= diluted crude supernatant) were constant and differences in OD_600_ at sampling time are normalized, the slope is a direct indicator for the amount of active enzyme produced by the according strain. The data used for calculation of the slope consists of four hours of kinetic measurement. Plotted values are averages of n = 3 biological replicates with error bars showing standard deviation.

### 4.3 Determining *gfp* expression levels on single-cell (flow cytometer) and population (plate reader) base

To test whether our system has an influence on *gfp* gene expression, two different approaches were made. First, it was checked on a single-cell level if more copies of *gfp* genes equal higher amounts of GFP protein and what the share of expressing cells was compared to the total cell population.

In general, observed fluorescence in the FL-1 channel appeared to positively correlate with cultivation time. However, for BsFLN051 with additional, plasmid-based *gfp* expression, values were consistently higher than for BsFLN045 (see Figure 2B).

Similarly, the share of fluorescent cells is significantly higher for the plasmid carrying strain BsFLN051 strain for all time points except t =11 h (see Figure 2C). At the same time, the number of non-fluorescent cells appears to be significantly lower if the plasmid is present, especially in early growth phases. To further verify above-mentioned results, a population-based fluorescence measurement was done in microtiter plates with the flow cytometry samples (see Supplemental Material Figure S1).

### 4.4 Application of CopySwitch to change *mPLC* expression levels

After showing the principle of CopySwitch with the easy-to-monitor GFP reporter protein, we applied the system to the expression of a secreted and industrially relevant enzyme. The phospholipase mPLC from *B. cereus* SBUG 516 has been chosen, which can be applied for the degumming of oils (Elena et al., 2017; Jiang et al., 2015). Furthermore, it already has been shown to be expressed in and secreted from *B. subtilis* and its activity can be assayed photometrically (Durban et al., 2007). The mPLC cleaves the substrate *p*-NPPC and thereby releases *p*-nitrophenol, which can be measured as increase of absorption at 410 nm. In total, four different time points were assayed (t6, t12, t24 and t48; number corresponds to hours after inoculation). Since every parameter was kept the same for every sample, the amount of enzyme in the sample directly correlates to the slope of the absorption change over time. OD_600_ differed slightly between triplicates and strains, but OD_410_ values were later normalized to this for each biological replicate before calculations. To make sure the increase of absorption is not due to background activity from the cultivation supernatant, each sample was additionally measured without substrate. One strain was included as a negative control, which contained no *mPLC* gene at all (BsMKA005). Furthermore, to exclude autohydrolysis of the substrate, only buffer containing substrate was measured. After four hours of measurement at 30°C, no significant increase of absorption could be detected for a) the negative control strain with and without substrate, b) buffer with and without substrate and c) mPLC-containing strains without substrate. All these controls strongly suggest mPLC as the sole source of *p*-NPPC-cleaving activity. After six hours of cultivation, almost no enzymatic activity can be detected in the diluted supernatant used for the assay, indicating expression has not started yet. After 12 and 24 hours of cultivation, the strain containing plasmid and chromosomal copies (BsFLN066) yields an about 1.3-fold increase of the slope (= activity) over the strain containing only a chromosomal *mPLC* copy (BsFLN064) in both cases. After 48 hours the activity difference drops to about 1.1-fold, still favoring BsFLN066 (see Figure 3B).

## 5. Discussion

For CopySwitch being applicable as a tool for the *B. subtilis* community, we first had to find the right transformation parameters. Our results suggest these to be approximately 2000 bp for each homology region, a regeneration time of four hours and a kanamycin concentration of 25 μg/ml. The overall increased values, compared to standard chromosomal integration (500 bp, one hour, 12.5 μg/ml) may stem from the plasmid nature of the CopySwitch product, which requires separate replication machinery and replication of a circularized plasmid. The *ori* used in our system has a reported copy number of six units/chromosome (Titok et al., 2003). Since the used antibiotic resistance marker against kanamycin is also increased in gene copy number, this may explain the possibility of using higher antibiotic concentrations. In addition, we tried using the backbone of the high copy number plasmid pMSE3 (Silbersack et al., 2006) (data not shown). These attempts did not show robust and reproducible results, likely due to the necessity of having four kb of adjacent chromosomal DNA for efficient homologous recombination on the plasmid, resulting in the increase of copy numbers of ORFs in these regions. For the *pksX* locus used in this work, this implies a copy number increase of *ymzD, ymcC* (both putative integral inner membrane proteins), *pksS* (cytochrome P450 of bacillaene metabolism) and *ymzB* (hypothetical protein) to several hundreds (Silbersack et al., 2006), posing the risk of disadvantageous effects. This locus had been chosen, because it contains a secondary metabolite cluster for the polyketide bacillaene (Butcher et al., 2007), which has proven to not be essential under laboratory conditions in our previous experiments (data not shown). Deletion of this cluster furthermore reduces the genome by 76.5 kb, which means less metabolic cost for replication and unnecessary protein production. It has already been shown elsewhere, that efficiency of recombinational transfer may be region dependent (Tomita et al., 2004a). The integration of completely synthetic and unique regions into the chromosome could be a way to address this issue. To find out more about the metabolic stress applied by additional gene copy numbers, growth curves were recorded. There were no visible differences neither in lag-, nor in log-phase, indicating that the additional burden of plasmid replication in BsFLN051 is not interfering with growth (see Figure 2A). However, in the transition stage from log- to stationary phase, BsFLN051 seems to have a slight disadvantage and also does not reach the final optical density of BsFLN045. This is likely due to the nature of the acetoin-inducible promoter, which is controlled by catabolite repression, meaning expression from it only starts once residual glucose in the medium is exhausted. As shown elsewhere, the growth is impaired from the onset of protein expression. This effect is getting more pronounced as the number of promoter-gene constructs rises (Silbersack et al., 2006). With our second test system (mPLC), those effects were not observed, probably due to a lower cultivation temperature of 30°C and the omission of glucose.

Since the system is intended for fast optimization of heterologous gene expression, we performed test expressions of *GFPmut2* with (BsFLN051) and without (BsFLN045) CopySwitch. The read-out on a single-cell level indicates not only higher fluorescence of the plasmid-bearing strain, but also an increased share of fluorescent cells. Apparently, additional *gfp* transcription facilitated by CopySwitch plasmid p14073 appears to increase the total amount of GFP in cells. The reduction of non-fluorescent cell population in BsFLN051 might be due to stronger deregulation of the utilized *acoA* promoter through titration of the CcpA (Carbon catabolite control protein A) repressor proteins by the increased copy number of CcpA binding sites in comparison to the single chromosomal promoter. Since glucose is contained in the medium to ensure promoter repression and therefore undisturbed growth before the onset of protein production, *ccpA* plays an important role in our system (Ali et al., 2001). Additionally, a population-based photometric measurement was conducted in microtiter plates directly after flow cytometry. As expected, the fluorescence intensity (normalized by OD_600_) is following the same pattern, rising over time and being higher in the plasmid-bearing BsFLN051 (see Supplemental Material Figure S1).

To strengthen our hypothesis, another experiment was performed with strains containing either no *mPLC* gene (BsMKA005), one *mPLC* gene integrated into the chromosome (BsFLN064) or *mPLC* both in the chromosome and on a low copy number plasmid (BsFLN066). The slope of the enzymatic activity (= absorption increase at 410 nm) was taken as read-out for comparison of the strains, after having been normalized against their respective OD_600_ values at the time of sampling. BsMKA005 showing no activity neither with nor without chromogenic substrate, suggests that mPLC is the enzyme responsible for absorption change in the supernatants of the remaining two strains. Since the negative control of buffer plus substrate also did not show any increase of absorption over time, autohydrolysis seems to play an insignificant role. After a cultivation time of six hours, corresponding to the end of exponential phase, the first samples were taken. A very shallow slope indicates almost no activity for all three strains, which is in accordance to previous publications, showing maximum induction of acetoin catabolism after exponential phase (Ali et al., 2001). As expected, after 12 and 24 hours, both BsFLN064 and BsFLN066 had increased levels of activity compared to BsMKA005, with BsFLN066 outcompeting BsFLN064 by 1.3-fold each time. Since activity values are OD_600_-normalized and every parameter despite copy number has been kept constant, it is likely that CopySwitch is leading to the higher amount of enzyme in the supernatant, therefore being responsible for higher activity. This effect is still visible after 48 hours of cultivation, although dropping down to 1.1-fold in favor of BsFLN066.

## 6. Conclusions

In comparison to other means of plasmid transformation in *B. subtilis*, CopySwitch has the great advantage of using naturally occurring competency. Other methods require either special *E. coli* strains for plasmid propagation, special devices for electroporation or tedious preparation of very fragile protoplasts. Additionally, all of the above-mentioned methods are either not well suited for high-throughput applications due to lack of robustness or require expensive, specialized equipment. The CopySwitch method is as easy as growing *B. subtilis* in a competence-inducing medium, adding a linearized plasmid and expression mix and, after a short incubation period, plating on agar-plates. All these steps can be automated and parallelized with standard liquid-handling platforms.

In summary, our method comprises a very easy and straightforward tool for tuning of gene expression, which is scalable and easily automatizable. Future applications beyond the scope of this work could be an easy tuning of metabolic pathways by performing CopySwitch with complete pathways or subsets of these.

## 7. Declarations

### 7.1 Acknowledgements

We like to thank Alex Elsholz for providing pYC121 as a template for GFPmut2, Dr. Zeigler for providing BsKO7 and Dr. Boettcher for providing p14094 as a template for mPLC.

### 7.2 Author contributions statement

FN conducted the experiments. FN, FB and JK analyzed the data, and prepared the manuscript. All authors read and approved the final manuscript.

### 7.4 Availability of data and materials

The flow cytometry datasets analyzed during the current study are available from the corresponding author on reasonable request.

